# Aging Converts Microglial Repopulation into Maladaptive Reprogramming that Exacerbates Cognitive Deficits

**DOI:** 10.64898/2026.01.08.698519

**Authors:** Rui Yamada, Hirotaka Nagai, Yunhui Zhu, Chisato Numa, Masayuki Taniguchi, Tomoyuki Furuyashiki

## Abstract

**Background and Purpose:** Aging has been associated with neuroinflammation and cognitive decline. Microglial repopulation after pharmacological depletion has been proposed as a strategy to alleviate microglia-driven neuropathology. However, the effects of microglial repopulation on age-related cognitive decline remain largely unexplored. In the present study, we examined how microglial repopulation affects the transcriptomic profiles of cortical microglia and the decline in prefrontal cortex-dependent cognitive function in aged mice.

**Experimental Approach:** Young and aged male C57BL/6J mice were used in this study. Microglial depletion was induced by treatment with PLX3397, a CSF1R inhibitor, followed by microglial repopulation after drug withdrawal. Microglia isolated from the entire cerebral cortex were subjected to bulk RNA-sequencing analysis. The visual discrimination test followed by the response direction test was conducted to assess sensory learning and attentional set shifting abilities, respectively.

**Key Results:** Repopulated cortical microglia in aged, but not young, mice exhibited aberrant gene expression patterns, including reduced expression of microglial identity genes, reprogramming of innate and adaptive immune-related gene expression, and derepression of neuronal gene expression. Furthermore, microglial repopulation selectively impaired visual discrimination learning and attentional set shifting in aged mice.

**Conclusion and Implications:** These findings demonstrate that aging converts microglial repopulation into maladaptive reprogramming that exacerbates cognitive decline, possibly through aberrant gene expression programs. Therefore, the therapeutic potential of this approach must be carefully evaluated with respect to specific behavioral domains and disease contexts, including aging.

## 1. Introduction

The global population is aging rapidly, and aging itself represents a major risk factor for cognitive decline and dementia(Prince et al., 2016; Prince et al., 2013). A growing body of evidence indicates that neuroinflammation plays a central role in this process, with microglia serving as key mediators(Salter & Stevens, 2017). Transcriptomic and epigenetic analyses have revealed extensive age-dependent remodeling of microglial gene expression(Li et al., 2023). In aged mice, microglia exhibit reduced phagocytic capacity, leading to the accumulation of engulfed cellular debris, such as myelin, within lysosomes(Safaiyan et al., 2021). Moreover, distinct microglial subtypes have been identified in the aged brain, including lipid droplet-accumulating microglia, which are characterized by heightened inflammatory gene expression, impaired phagocytosis, and neurotoxic properties(Marschallinger et al., 2020). Microglial activation is also a prominent feature in age-related neurodegenerative conditions, such as Alzheimer’s disease, in which neuroinflammation is thought to exacerbate neuropathological progression(Leng & Edison, 2021).

To investigate the functional contribution of microglia to brain aging and disease, pharmacological depletion of microglia followed by their repopulation has been widely employed(Han, Zhu, Zhang, & Harris, 2019). Microglia originate from yolk sac progenitors and colonize the brain during early development, thereafter maintaining their population through self-renewal throughout adulthood(Ginhoux et al., 2010). Their survival critically depends on signaling through the colony stimulating factor 1 receptor (CSF1R) and its ligands, IL-34 or CSF-1, in a brain region-specific manner(Kana et al., 2019; Y. Wang et al., 2012). Consistent with this, treatment with CSF1R inhibitors, such as PLX3397, eliminates nearly the entire microglial population within weeks(M. R. Elmore et al., 2014). Upon withdrawal of the inhibitor, microglia rapidly repopulate the brain, likely derived from residual surviving microglia(Huang et al., 2018). Importantly, microglial repopulation has been proposed to reset maladaptive microglial states that emerge in various disease contexts, thereby alleviating microglia-driven neuropathology. Indeed, previous rodent studies have reported that microglial repopulation mitigates behavioral and cognitive deficits associated with aging, Alzheimer’s disease models, and physical insults, such as radiation and traumatic injury (Colella et al., 2024; M. R. P. Elmore et al., 2018; Han et al., 2019; Hu et al., 2025; W. Wang et al., 2023).

However, microglial depletion and repopulation are not universally beneficial. Microglial depletion may worsen cognitive function, may variably ameliorate chronic stress-induced behavioral alterations, and may even abolish the therapeutic effects of electroconvulsive stimulation(Rimmerman et al., 2022; Yegla, Boles, Kumar, & Foster, 2021). Microglial repopulation has been reported to reduce hippocampal synaptic density under basal conditions and to induce a relapse of chronic stress-induced behavioral deficits that had previously been mitigated with microglial depletion(Weber et al., 2019; Wickel et al., 2024). These observations suggest that the consequences of microglial depletion and repopulation are highly context-dependent, influenced by the pre-existing microglial state as well as the physiological or pathological condition of the brain. Therefore, the therapeutic potential of this approach must be carefully evaluated with respect to specific behavioral domains and disease contexts. In this regard, the effects of microglial repopulation on age-related cognitive decline remain largely unexplored, especially cognitive deficits associated with impaired prefrontal function(Morrison & Baxter, 2012; H. Nagai, 2024).

In the present study, we examined how microglial depletion followed by repopulation affects the transcriptomic profiles of cortical microglia and the decline in prefrontal cortex-dependent cognitive function in aged mice. We found that repopulated cortical microglia in aged, but not young, mice exhibited aberrant gene expression patterns, including reduced expression of microglial identity genes, reprogramming of innate and adaptive immune-related gene expression, and derepression of neuronal gene expression. Furthermore, microglial repopulation selectively impaired learning and cognitive flexibility in aged mice, as assessed by visual discrimination and response direction tests. Together, these findings demonstrate that microglial repopulation drives age-specific gene expression programs that exacerbate, rather than ameliorate, age-related cognitive decline.

## 2. Methods

### 2.1. Animals

Male C57BL/6J mice, either young (4-8 weeks old) or aged (67-84 weeks old), were obtained from Jackson Laboratories Japan (Yokohama, Japan). After arrival, mice were group-housed (4-5 per cage) for at least one week to acclimate and were maintained in a temperature- and humidity-controlled facility under a 12-h light/dark cycle, with ad libitum access to food and water. For microglial depletion and repopulation, mice were fed AIN-76A chow containing PLX3397, a CSF1R inhibitor, at a concentration of 0.029% (w/w) (Research Diets, New Brunswich, NJ, USA) for four weeks to deplete microglia, followed by an additional four-week withdrawal period to allow microglial repopulation. For behavioral testing, mice were individually housed beforehand. At the time of behavioral testing, young mice were 8-12 weeks old and aged mice were 75-92 weeks old.

All experimental procedures were designed to minimize animal suffering and to reduce the number of animals used, in accordance with the principles of the 3Rs (Replacement, Reduction, and Refinement). Animal health and welfare were monitored daily, and humane endpoints were applied when necessary. All procedures were conducted in accordance with the ARRIVE guidelines and the NIH Guide for the Care and Use of Laboratory Animals and were approved by the Animal Care and Use Committees of Kobe University Graduate School of Medicine.

### 2.2. Immunostaining

Immunostaining was performed as previously described(M. Nagai, Nagai, Numa, & Furuyashiki, 2020; Numa et al., 2019). Briefly, mice were anesthetized by intraperitoneal injection of sodium pentobarbital (100 mg/kg; Nacalai Tesque, Kyoto, Japan) and perfused transcardially with saline, followed by 4% paraformaldehyde in 0.1 M sodium phosphate buffer. Brains were removed and post-fixed in the same fixative overnight at 4°C. Coronal brain sections (40 μm thick) were cut using a vibratome and stored in cryoprotectant until further processing.

For immunostaining, free-floating sections were incubated for 1 h at room temperature (RT) in blocking solution consisting of Dulbecco’s modified phosphate buffered saline (D-PBS, 07269-84, Nacalai Tesque Inc., Kyoto, Japan) supplemented with 1% normal donkey serum (017-000-121, Jackson ImmunoResearch, West Grove, PA, USA) and 0.3% Triton X-100. Sections were then incubated overnight at 4°C with guinea pig anti-TMEM119 (400 004, Synaptic Systems; 1:500) as the primary antibody diluted in the same blocking solution. After three washes in D-PBS containing 0.3% Triton X-100 at RT, sections were incubated for 2 h at RT with Alexa Fluor 647-conjugated anti-guinea pig IgG (706-605-148, Jackson ImmunoResearch; 1:1000) as the secondary antibody. After three additional washes, sections were stained with Hoechst 33342 (Thermo Fisher Scientific, Waltham, MA, USA; 1:5000 in D-PBS) for 20 min at RT, and then washed twice in D-PBS.

Stained sections were mounted on glass slides (Matsunami Glass, Kishiwada, Japan) and coverslipped with ProLong Gold Antifade Mountant (Thermo Fisher Scientific). Images were acquired through a 10x objective lens on an LSM700 confocal microscopy (Carl Zeiss Microscopy GmbH, Jena, Germany). For image analysis, fluorescent signals were quantified by ImageJ.

### 2.3. Behavioral tests

Prior to behavioral testing, mice were food-restricted to approximately 80% of their baseline body weight. All tests were conducted in an operant chamber equipped with a touchscreen display and an integrated food tray connected to an automated pellet dispenser (O’Hara & Co., Ltd., Tokyo, Japan).

During the habituation phase, on day 1, mice were introduced to the chamber and allowed to obtain five food pellets (1811213, TestDiet, Richmond, IN, USA, 10 mg each) manually placed in the tray. On day 2, mice were given one food pellet delivered to the tray every 10 s for 15 min (approximately 90 pellets) via the automated food dispenser. On subsequent days, mice were presented with visual stimuli (horizontal and vertical lines) displayed simultaneously on two windows of the touchscreen interface and were required to touch either window to receive a food pellet. Each session lasted up to 1 h or until 150 touches were recorded, whichever occurred first. Training was repeated on consecutive days until they made at least 30 touches in a session.

Following habituation, mice underwent a visual discrimination test in which they were required to distinguish between horizontal and vertical line stimuli to obtain a food pellet. Pellets were delivered to the tray only when mice touched the stimulus (horizontal or vertical lines) designated as correct throughout the experiment. Each session lasted up to 1 h or until the predefined number of trials was completed. Control mice performed up to 100 trials per session in two sessions per day, whereas mice after microglial repopulation performed up to 150 trials per session in one session per day. Training continued daily until mice achieved ≥80% correct responses within the first 100 trials of a session or ≥75% accuracy across two sessions (either consecutive or non-consecutive). The task was conducted for up to 12 days in control mice and up to 16 days in mice after microglial repopulation.

After completing the visual discrimination test, mice proceeded to the response direction test. In this test, mice were required to touch the touchscreen window designed as correct (either left or right side) to receive a food pellet, regardless of the visual stimulus presented. Each session lasted up to 1 h or 100 trials. The mice completed one session per day for five consecutive days.

A subset of the behavioral data with control mice are reported elsewhere(Yamada et al., 2026).

### 2.4. RNA-seq Analysis of Isolated Microglia

RNA sequencing (RNA-seq) analysis was performed as previously described(Horikawa et al., 2024; Mishima, Taniguchi, Matsushita, Tian, & Furuyashiki, 2023). Mice were deeply anesthetized with sodium pentobarbital (100 mg/kg, intraperitoneal) and perfused transcardially with calcium- and magnesium-free Hank’s Balanced Salt Solution (HBSS). Brains were removed, and the cerebral cortices were dissected in 1 × HBSS. After replacing the buffer with HBSS containing RNase inhibitors (HBSS-Ri), the tissue was homogenized using a Dounce homogenizer and passed through a 40-µm strainer.

The homogenate was centrifuged at 800 × g for 5 min at 4°C, and the cell pellet was resuspended in 30% Percoll (prepared by diluting 60% Percoll 1:1 with HBSS-Ri). Th suspension was centrifuged at 800 × g for 25 min at 4°C, and the myelin-containing supernatant was carefully removed. The resulting cell pellet was washed and resuspended in MACS buffer (19:1 mixture of AutoMACS Rinsing Solution and MACS BSA Stock; Miltenyi Biotec GmbH, Bergisch Gladbach, FRG). CD11b-positive cells, predominantly microglia in the brain, were isolated using CD11b MicroBeads (130-126-725; Miltenyi Biotec) with the autoMACS Pro Separator (Miltenyi Biotec), according to the manufacturer’s protocol. The isolated microglia were centrifuged at 2,000 × g for 10 min at 4°C, washed with PBS containing RNase inhibitors, centrifuged again, and the final pellet was stored at −80°C until RNA extraction.

Total RNA was extracted using the RNAdvance Cell kit (Beckman Coulter, Brea, CA, USA), according to the manufacturer’s protocol. RNA quality was assessed using a Bioanalyzer 2100 (Agilent Technologies, Santa Clara, CA, USA) with the RNA Pico chip. RNA yield and integrity were as follows: 12,115.6 ± 679.5 pg/μL and RIN 7.8 ± 0.1 (young control), 7,884 ± 660.5 pg/μL and RIN 7.8 ± 0.1 (young repopulation), 9,330.8 ± 404.2 pg/μL and RIN 7.9 ± 0.1 (aged control), 622.4 ± 58.3 pg/μL and RIN 7.6 ± 0.2 (aged depletion), and 9,581.8 ± 399.0 pg/μL and RIN 7.8 ± 0.1 (aged repopulation). Complementary DNA was synthesized using the SMART-Seq v4 kit (Takara Bio Inc., Kusatsu, Shiga, Japan), and sequencing libraries were prepared with the Nextera XT kit (Illumina, San Diego, CA, USA). Sequencing was performed on a NovaSeq X Plus platform at Novogene Japan (Tokyo, Japan).

Transcript quantification was conducted using Kallisto (v0.46.1) with Ensembl GRCm38 mouse transcript references, as previously described(Horikawa et al., 2024). Differential expression analysis was performed at the transcript level using Sleuth, and TPM (transcripts per million) values for all transcript isoforms were then summed to obtain gene-level expression values for downstream analyses. Of 35,845 annotated genes, 13,222 were consistently detected across all samples and were included in downstream analyses. Z-scores of gene expression were calculated, and hierarchical clustering was applied for pattern analysis.

Gene ontology (GO) enrichment analysis was performed using the publicly available Gene Ontology resource (https://geneontology.org/). Subsets of genes showing statistically significant differential expression (p < 0.05) identified in each analysis were analyzed. GO terms containing more than three and fewer than 500 assigned genes were included. Up to five out of the top ten GO terms, ranked by fold enrichment, were displayed. To assess synapse-related gene expression, SynGO analysis was performed(Koopmans et al., 2019). Because SynGO is based on human gene annotations, mouse genes were first converted to their human orthologs. Of the 13,222 detected mouse genes, 13,075 were successfully converted, and 1,149 genes (8.8%) were annotated as synapse-related in the SynGO database.

### 2.5. Statistical analyses

All data are presented as means ± SEM. Statistical analyses were performed using Prism version 10.5 (GraphPad Software, San Diego, CA, USA). A p value of less than 0.05 was considered statistically significant. Comparisons between two groups were conducted using a two-tailed unpaired Student’s *t*-test (Figs. 3A-F,5A,6C, and 6D). Comparisons among three or more groups were performed using one-way ANOVA followed by Holm-Šidak’s multiple comparisons test (Figs. 1C,2B,2F,3H,4D,5F and 6F). One-sample *t*-tests were used to determine whether the observed values significantly exceeded the chance level (50%) (Fig. 6F).

## Results

### Repopulated cortical microglia show age-specific aberrant gene expression programs

To determine whether microglial repopulation following depletion can alleviate age-related changes in gene expression and rejuvenate cortical microglia, we performed transcriptomic analyses of cortical microglia from young and aged mice before and after microglial repopulation. Microglia were depleted by treating mice with PLX3397, a CSF1R inhibitor, for four weeks, followed by a four-week withdrawal period to allow microglial repopulation (Fig. 1A). Immunostaining for the microglial marker TMEM119 confirmed that this treatment effectively depleted microglia in both young and aged mice, although depletion was less complete in aged mice than in young mice, and that microglia subsequently repopulated the cerebral cortex to approximately 70% of baseline levels (Fig. 1B,C).

**Figure 1.**
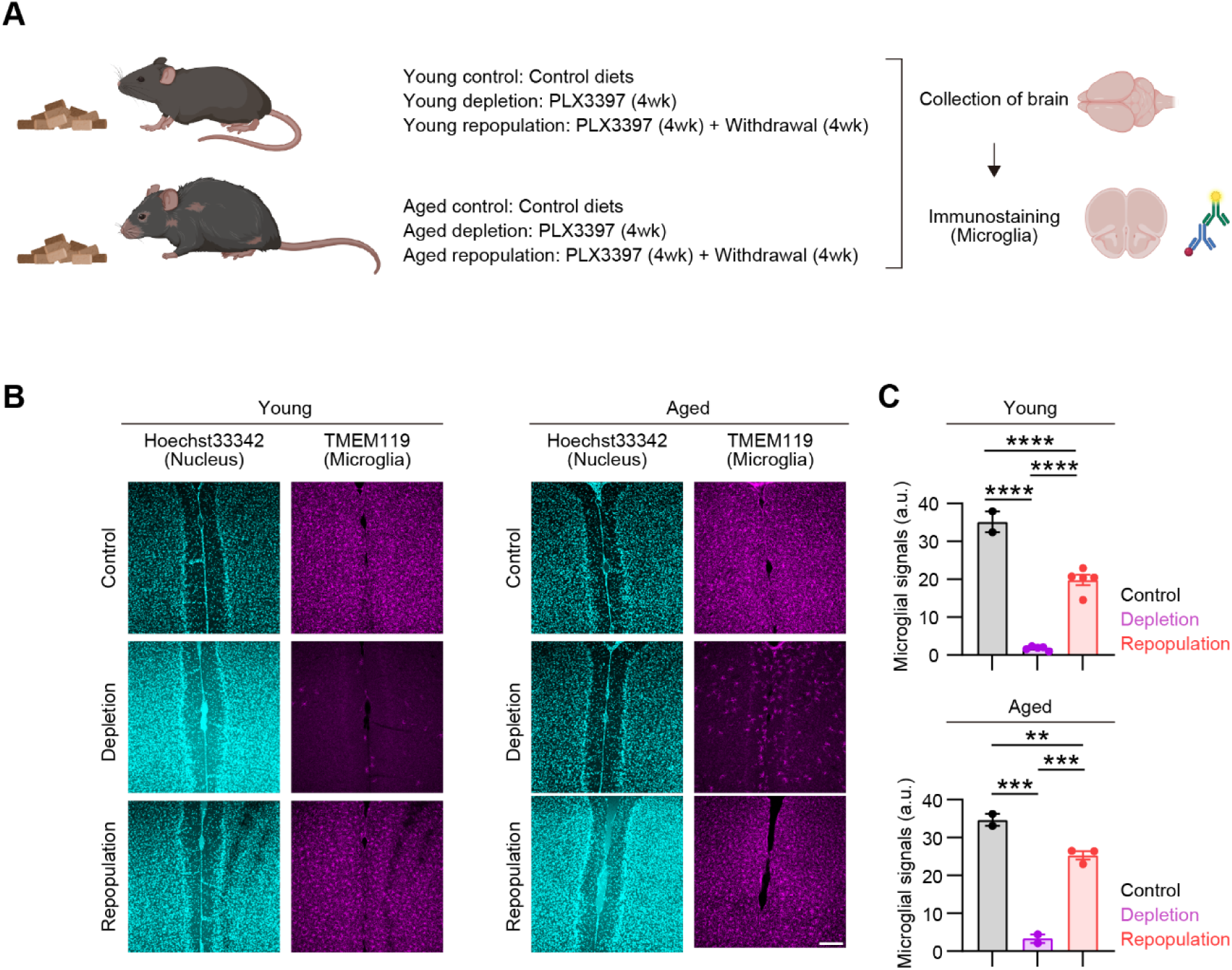
Microglial depletion and repopulation following PLX3397 treatment. (A) Experimental design for microglial depletion and repopulation. Young and aged mice were fed control diets (control) or PLX3397-containing diets for 4 weeks, either without (depletion) or with (repopulation) an additional 4-week withdrawal period. Brains were collected for immunostaining. (B) Immunostaining of microglial markers (TMEM119) with nuclear counterstaining (Hoechst 33342). Scale bar = 200 μm. (C) Signal intensity of TMEM119 immunoreactivity. Values are expressed as means ± SEM. *p < 0.05, ***p < 0.001, ****p < 0.0001 for Holm-Sidak’s multiple comparisons test following one-way ANOVA.

We then isolated cortical microglia from the cerebral cortex using CD11b-based magnetic bead purification from young mice without depletion (young control) or after repopulation (young repopulation), and from aged mice without depletion (aged control), after depletion (aged depletion), or after repopulation (aged repopulation) (Fig. 2A). In young mice subjected to microglial depletion, insufficient numbers of cortical microglia could be obtained because PLX3397 treatment resulted in near-complete depletion. We then analyzed gene expression patterns of isolated cortical microglia using bulk RNA-seq analysis. We first examined the expression of canonical microglial markers. Whereas *P2ry12* expression was similar across experimental conditions, microglial repopulation in most aged mice (6/8), but not in young mice, reduced the expression of other microglial markers (*Hexb*, *Cx3cr1*, *Tmem119*, *Sall1*, *Fcrl2*) to negligible levels (Fig. 2B), indicating that microglial repopulation in aged mice alters fundamental microglial properties.

**Figure 2.**
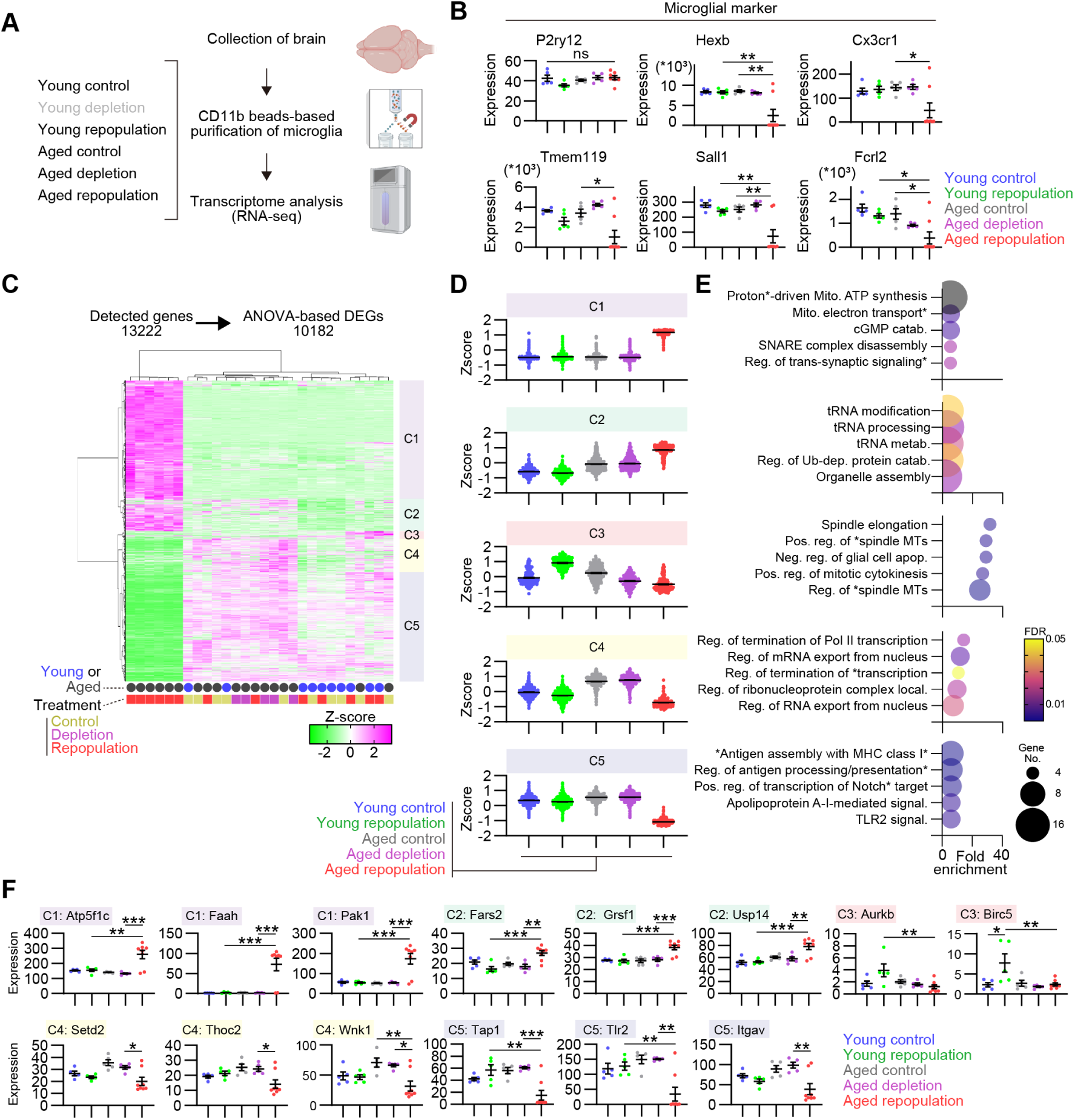
Repopulated cortical microglia show age-specific aberrant gene expression programs. (A) Experimental design for RNA-seq analysis of isolated microglia. Young and aged mice were fed control diets (control) or PLX3397-containing diets for 4 weeks, either without (depletion) or with (repopulation) an additional 4-week withdrawal period. Cortical microglia were isolated using CD11b magnetic beads for RNA-seq analysis. (B) Expression levels of canonical microglial marker genes, expressed as transcripts per million (TPM). (C) Heatmap and hierarchical clustering of ANOVA-based differentially expressed genes (DEGs). Animal age (young, blue; aged, dark gray) and treatment condition (control, ocher; depletion, magenta; repopulation, red) are indicated below the heatmap. Five gene clusters (C1-C5) were defined. (D) Z-scored expression levels of each gene were averaged across individuals within each group for each cluster. (E) Gene ontology enrichment analysis of DEGs within each cluster. The false discovery rate (FDR) and the number of included genes for each GO term are indicated by color and circle size, respectively. (F) Expression levels of representative genes from each cluster, expressed as TPM. Data are presented as mean ± SEM. *p < 0.05, **p < 0.01, ***p < 0.001; ns, not significant for Holm-Sidak’s multiple-comparisons test following one-way ANOVA.

ANOVA across all experimental groups identified 10,182 differentially expressed genes (DEGs). Hierarchical clustering of these DEGs revealed five major gene clusters exhibiting distinct expression patterns (Fig. 2C). Among these DEGs, many genes were upregulated or downregulated after microglial repopulation only in aged mice, without apparent age-related changes, forming the largest clusters, C1 and C5, respectively (4,005 and 3,712 genes) (Fig. 2D). Fewer genes showed age-associated upregulation and were either further increased or decreased after microglial repopulation only in aged mice, comprising two distinct clusters, C2 and C4, respectively (1,086 and 1,123 genes). A minor but significant subset of genes exhibited increased expression in young mice and decreased expression in aged mice after microglial repopulation, as represented by the smallest cluster, C3 (256 genes).

Gene Ontology (GO) enrichment analyses revealed that genes within these clusters, which display distinct expression patterns with aging and/or microglial repopulation, are closely linked to mechanisms regulating microglial states and functional responses (Fig. 2E). Specifically, C1 genes are predominantly associated with mitochondrial oxidative phosphorylation/electron transport, with a smaller subset related to synaptic functions. This cluster includes genes involved in multiple components of the mitochondrial electron transport chain and ATP synthase (e.g., *Nduf*, *Sdh*, *Uqcr*, *Atp5*), as well as genes involved in the cannabinoid pathway (e.g., *Faah*, *Cnrip1*), cytoskeletal and regulatory kinases (e.g., *Pak1*), and cyclic nucleotide signaling modulators (e.g., *Pde* family genes) (Fig. 2F). C2 genes are mainly associated with tRNA-related processes, ubiquitin-dependent protein catabolic processes, and organelle assembly. This cluster contains genes encoding mitochondrial RNA-binding proteins and aminoacyl-tRNA synthetases (e.g., *Grsf1*, *Fars2*, *Tars2*), tRNA modification enzymes (e.g., *Dalrd3*, *Trmt10b*, *Pus7*, *Elp2*), autophagy- and metabolic signaling components (e.g., *Pik3c3*, *Rheb*), and regulators of ubiquitin-proteasome-mediated protein turnover (e.g., *Usp14*, *Uchl5*, *Ube3a*, *Zfand2a*). C3 genes are primarily associated with cell cycle progression and mitotic regulation. This cluster includes genes involved in cell cycle and mitotic control (e.g., *Kif11*, *Aurkb*, *Birc5*, *Cdca8*, *Ccnb1*, *Rad21*, *Ect2*), as well as genes related to growth factor signaling and cell-cell interactions (e.g., *Gas6*, *Igf1*, *Sdk1*). C4 genes are mainly associated with transcriptional and post-transcriptional regulation. This cluster contains genes involved in transcriptional regulation and chromatin modification (e.g., Setd2, Setx, Ppp1r10), RNA processing and mRNA export (e.g., *Scaf4*, *Thoc2*, Iws1), and nuclear structure and signaling scaffolding (e.g., *Tpr*, *Akap8l*, *Wnk1*). C5 genes are predominantly associated with antigen processing and presentation and immune receptor signaling. This cluster includes genes involved in major histocompatibility complex (MHC) class I antigen processing and peptide loading (e.g., *Tap1*, *Tap2*, *Tapbpl*, *Calr*, *Pdia3*, *Erap1*, *B2m*, *H2-Oa*, *H2-Ob*, *Hfe*), regulation of Notch signaling and transcriptional control (e.g., *Notch1*, *Maml1*, *Maml2*, *Rbpj*, *Rbm15*), innate immune and inflammatory signaling (e.g., *Tlr2*, *Ripk2*, *Irak1*, *Tnip2*, *Pik3ap1*), and phagocytosis and cell adhesion (e.g., *Abca1*, *Abca7*, *Itgav*, *Itgb3*, *Rhoa*, *Mal3*).

Given that the largest gene clusters, C1 and C5, overall reflect gene expression changes specific to aged mice, microglial repopulation after aging gives rise to abnormal microglia with aberrant gene expression programs. These repopulated microglia likely adopt a dysfunctional state characterized by altered metabolic and phagocytic properties, along with a reduced capacity to respond to extracellular immune and inflammatory signals. Analysis of C2 and C4 clusters reveals that, although microglial repopulation rejuvenates a subset of age-related gene expression changes, particularly those associated with transcriptional and post-transcriptional regulation, it also exacerbates another subset, notably genes involved in protein translation and degradation. Moreover, C3 cluster indicates that, unlike in young mice, repopulated microglia in aged mice display reduced proliferative and mitotic capacity. Collectively, these results demonstrate that aging converts microglial repopulation into maladaptive reprogramming with aberrant gene expression patterns.

### Age-specific gene expression programs predominate over age-shared changes in repopulated microglia

To further delineate differences in gene expression among experimental conditions, we performed pairwise comparisons. Comparison of young repopulation and control mice identified DEGs associated with microglial repopulation in young mice (401 upregulated and 616 downregulated genes for young repopulation vs. young control), although the numbers were substantially lower than those observed with aging alone (1,516 upregulated and 406 downregulated genes for aged control vs. young control) (Fig. 3A,B). In contrast, the aged repopulation group differed markedly from all other groups, including the young repopulation group (4,700 upregulated and 3,007 downregulated genes for aged repopulation vs. young repopulation) (Fig. 3C), consistent with the clustering analysis described above. Notably, microglial depletion alone in aged mice induced much smaller changes (311 upregulated and 308 downregulated genes for aged depletion vs. aged control) (Fig. 3D). However, following repopulation, aged microglia acquired a markedly distinct profile, as evidenced by the large numbers of DEGs between aged depletion and repopulation groups (4,449 upregulated and 4,830 downregulated genes for aged repopulation vs. aged depletion), which were comparable to those observed between aged control and repopulation mice (4,322 upregulated and 4,729 downregulated genes for aged repopulation vs. aged control) (Fig. 3E,F). Thus, in aged mice, gene expression changes occurred predominantly during microglial repopulation after depletion.

**Figure 3.**
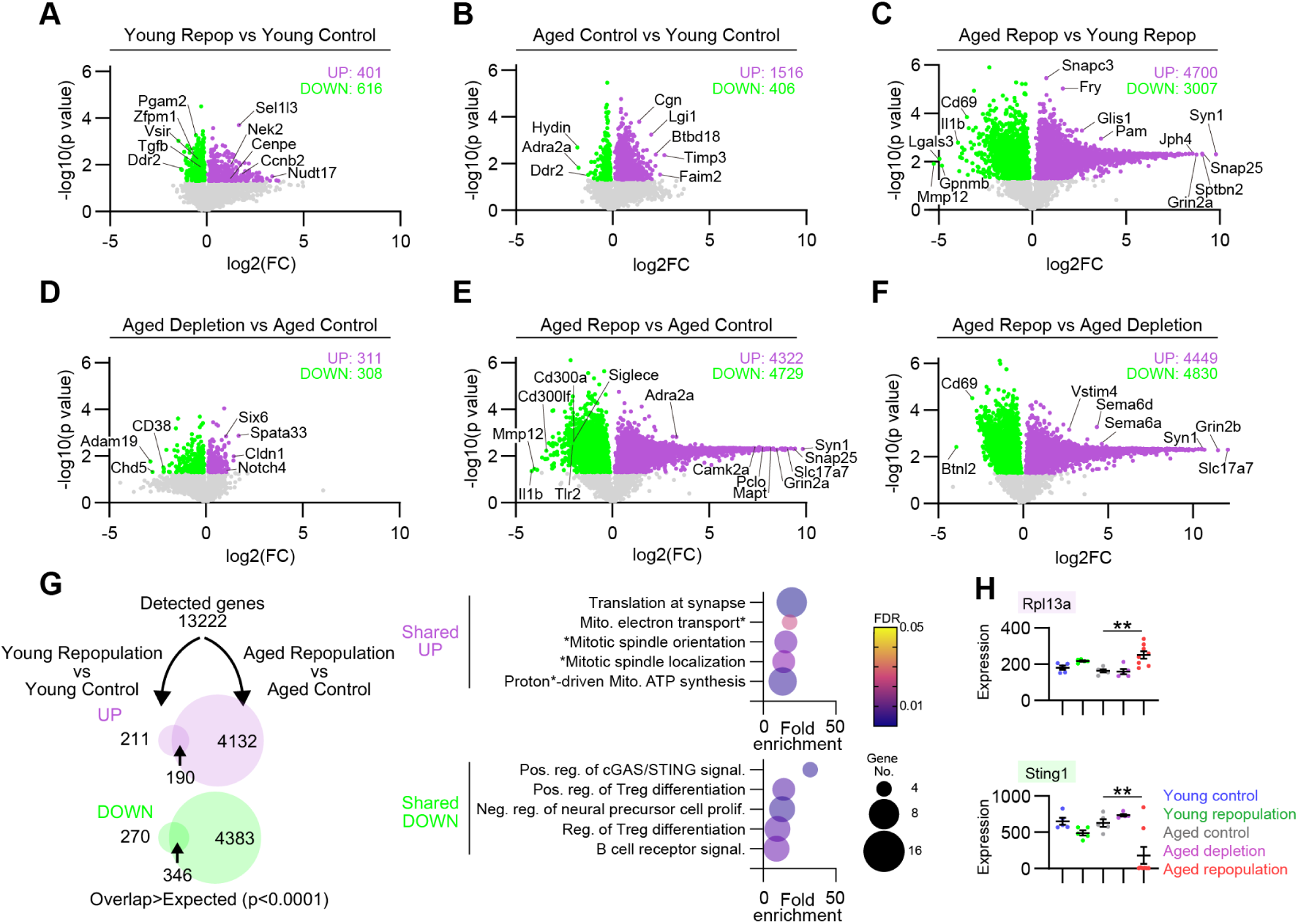
Age-specific gene expression programs predominate over age-shared changes in repopulated microglia. (A-F) Volcano plots comparing the indicated group pairs. The x-axis shows log_2_ fold change (FC) and the y-axis shows −log_10_ p values based on *t*-tests. Genes upregulated and downregulated with statistical significance are highlighted in magenta and green, respectively. (G) Venn diagrams depicting genes upregulated (magenta) and downregulated (green) in repopulated microglia relative to control microglia in young and aged mice (left and right circles, respectively). Gene ontology terms enriched for age-shared upregulated and downregulated genes are shown. The false discovery rate (FDR) and the number of included genes for each GO term are indicated by color and circle size, respectively. (H) Expression levels of representative age-shared upregulated and downregulated genes, expressed as transcripts per million. Data are presented as mean ± SEM. **p < 0.01 for Holm-Sidak’s multiple-comparisons test following one-way ANOVA.

Despite these age-specific effects, microglial repopulation still induced significantly overlapping changes in young and aged mice (Fig. 3G), indicating the presence of a shared transcriptional program, although this represents only a minor fraction of the changes observed in aged mice. GO enrichment analysis showed that commonly upregulated genes were enriched with ribosomal components for translation, components of the mitochondrial electron transport chain, and cell division-related processes, whereas commonly downregulated genes were enriched with immune signaling and differentiation-related pathways, such as the cGAS-STING and TGF-β signaling (Fig. 3H; see Fig. 4D for *Tgfb1* expression).

**Figure 4.**
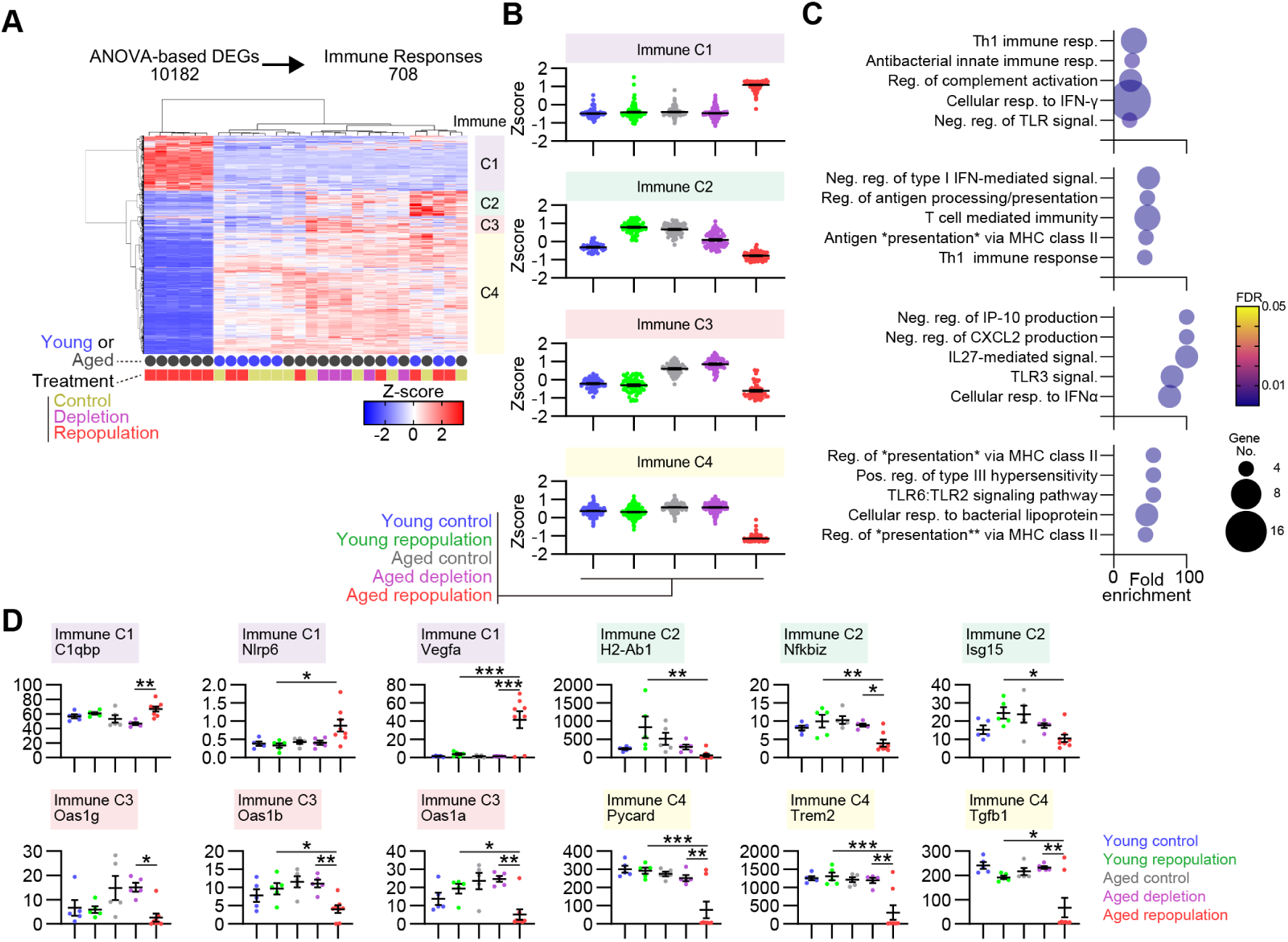
Microglial repopulation in aged mice reprograms innate and adaptive immune-related gene expression. (A) Heatmap and hierarchical clustering of immune response-related genes extracted from ANOVA-based differentially expressed genes (DEGs). Animal age (young, blue; aged, dark gray) and treatment condition (control, ocher; depletion, magenta; repopulation, red) are indicated below the heatmap. Four gene clusters (C1-C4) were defined. (B) Z-scored expression levels of each gene across individual within each group for each cluster. (C) Gene ontology enrichment analysis of genes within each cluster. The false discovery rate (FDR) and the number of included genes for each GO term are indicated by color and circle size, respectively. (D) Expression levels of representative genes from each cluster, expressed as transcripts per million. Data are presented as mean ± SEM. *p < 0.05, **p < 0.01, ***p < 0.001; ns, not significant for Holm-Sidak’s multiple-comparisons test following one-way ANOVA.

Collectively, these results demonstrate that age-specific gene expression programs that emerge during microglial repopulation predominate in repopulated microglia of aged mice, accompanied by minor yet significant changes shared across ages.

### Microglial repopulation in aged mice reprograms innate and adaptive immune-related gene expression

To further characterize immune-related gene expression changes across experimental conditions, we focused on genes annotated to immune response (GO:0006955) among the ANOVA-identified DEGs and subdivided them into four immune-related subclusters (immune C1-C4) based on their expression patterns (Fig. 4A). Most immune-related genes exhibited repopulation-dependent changes specifically in aged mice, forming the two largest clusters, immune C1 and C4, which were selectively upregulated (180 genes) and downregulated (391 genes), respectively, after repopulation in aged mice (Fig. 4B). In contrast, the smaller clusters, immune C2 and C3, both comprised genes that were upregulated after aging but downregulated after microglial repopulation (79 and 58 genes). However, their responses to repopulation in young mice differed between the clusters: immune C2 genes increased, whereas immune C3 genes remained unchanged.

GO enrichment analyses showed that immune C1 genes are enriched for Th1 immune responses, type II interferon signaling, and regulation of complement activation (Fig. 4C). This cluster includes genes involved in cell-cell interactions (*Sema4a*, *Vegfa*), innate immune and inflammasome-associated regulation (*Nlrp6*, *Sarm1*), complement- and immune-regulatory functions (*C1qbp*, *Cfp*, *Clu*, *Cd46*), as well as genes involved in diverse intracellular processes (Fig. 4D). Immune C2 genes are enriched for type I interferon signaling, antigen presentation, and Th1 immune responses. This cluster includes genes for transcriptional regulators (*Bcl3*, *Nfkbiz*), interferon-responsive genes (*Oas3*, *Adar*, *Isg15*), components of the antigen presentation machinery (*H2-Ab1*, *H2-Aa*, *H2-Eb1*, *Cd74*, *Nlrc5*), and immune effector or regulatory molecules (*Cst7*, *Fgl2*, *Serpinb9*, *Stx11*, *Cd8a*, *Rftn1*, *Slfn2*). Immune C3 genes consist predominantly of oligoadenylate synthetase family genes (*Oas1a*, *Oas1b*, *Oas1c*, *Oas1g*, *Oas2*) that mediate RNase L-mediated RNA degradation as a core antiviral defense mechanism, together with interferon-inducible nucleic acid sensors (*Ifi204*). Immune C4 genes are enriched for innate immune sensing and inflammatory signal transduction. This cluster includes genes associated with regulation of MHC class II pathways (*H2-Oa*, *H2-Ob*), microglial activation and phagocytosis (*Trem2*, *Fcgr1*, *Fcgr3*, *Fcer1g*, *Btk*), inflammasome assembly (*Pycard*), and Toll-like receptor signaling (*Tlr1*, *Tlr2*, *Tlr6*, *Cd14*, *Myd88*, *Tirap*, *Rela*).

Collectively, these findings demonstrate that microglial repopulation in aged mice drives aberrant, age-specific reprogramming of innate and adaptive immune-related gene expression.

### Microglial repopulation in aged mice derepresses neuronal gene expression

In the transcriptomic analyses described above, a subset of genes upregulated after microglial repopulation in aged mice was associated with synaptic functions (see cluster C1 in Fig. 2D,E). Consistently, several genes encoding presynaptic and postsynaptic proteins were robustly upregulated after microglial repopulation (Fig. 5A; also see Fig. 3E). Notably, GO enrichment analysis showed that the most highly upregulated genes (fold change greater than 100) were enriched for synaptic functions (Fig. 5B). Indeed, synapse-associated genes annotated in the SynGO database accounted for approximately half of these highly upregulated genes (Fig. 5C).

**Figure 5.**
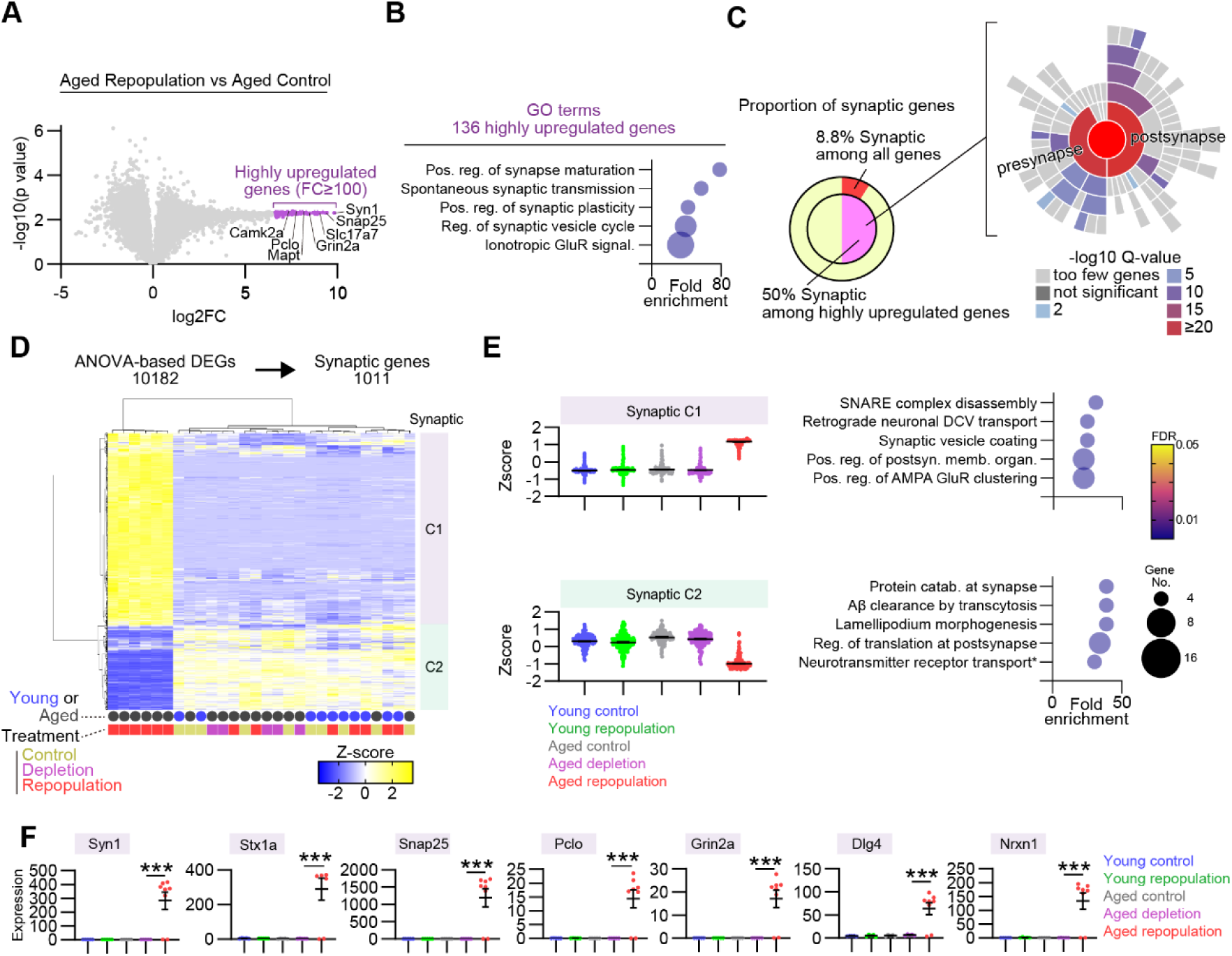
Microglial repopulation in aged mice derepresses neuronal gene expression. (A) Volcano plot comparing the aged repopulation and control groups. The x-axis shows log_2_ fold change (FC) and the y-axis shows −log_10_ p values based on *t*-tests. Highly upregulated genes (FC ≥ 100) with statical significance are highlighted in magenta. The same plot is shown in Fig. 2E with different genes highlighted. (B) Gene ontology (GO) terms enriched for the highly upregulated genes in the aged repopulation group. The false discovery rate (FDR) and the number of included genes for each GO term are indicated by color and circle size, respectively. (C) GO enrichment analysis of the highly upregulated genes in the aged repopulation group for synapse-related genes based on the SynGO database. Left: The proportions of synase-related genes among the highly upregulated genes or all genes are shown. Right: The q value for each GO term represented by each sector is indicated by color. (D) Heatmap and hierarchical clustering of synapse-related genes extracted from ANOVA-based differentially expressed genes (DEGs). Animal age (young, blue; aged, dark gray) and treatment condition (control, ocher; depletion, magenta; repopulation, red) are indicated below the heatmap. Two gene clusters (Synaptic C1 and C2) were defined. (E) Z-scored expression levels of each gene averaged across individuals within each group for each cluster. The FDR and the number of included genes for each GO term are indicated by color and circle size, respectively. (F) Expression levels of representative genes from each cluster, expressed as transcripts per million. Data are presented as mean ± SEM. ***p < 0.001 for Holm-Sidak’s multiple-comparisons test following one-way ANOVA.

To systematically characterize these changes, we extracted 1,011 synapse-associated, ANOVA-based DEGs using the SynGO database (Fig. 5D). Hierarchical clustering revealed two major expression patterns: synaptic C1 genes were upregulated, whereas synaptic C2 genes were downregulated, after microglial repopulation in aged, but not young, mice (Fig. 5E). GO enrichment analyses indicated that synaptic C1 genes were associated with synaptic organization, vesicle trafficking, and neuronal signaling pathways. This cluster includes genes encoding components of the presynaptic release machinery and vesicle recycling (*Syn1*, *Stx1a*, *Snap25*, *Vamp1/2*, *Syt1*, *Pclo*, *Rims2*, *Unc13a*), postsynaptic receptors and scaffolding proteins (*Grin2a/b*, *Gria2/3*, *Dlg4*, *Shank1*, *Homer1*), synapse-organizing adhesion molecules (*Nrxn1/2*, *Nlgn2/3*, *Cdh10*, *Pcdh10*, *Lrrtm2*, *Cntnap2*), and axon guidance molecules (*Epha4*, *Ephb2*, *Sema3a*, *Ntn1*) (Fig. 5F). Notably, transcripts encoding these synaptic and neuronal proteins were detected only in aged mice after microglial repopulation and were negligible in other conditions. Thus, these finding are unlikely to result from contamination of neuronal components in the isolated microglia.

In contrast, synaptic C2 genes were enriched for pathways regulating endocytic trafficking, membrane protein recycling, and cytoskeletal regulation. This cluster includes genes for clathrin-mediated endocytosis and endosomal sorting (*Cltc*, *Picalm*, *Ap1g1*, *Snx1*, *Vps35*, *Rab5b*, *Rab7*, *Rab11a*), actin regulatory modules (*Wasf2*, *Abi1*, *Cyfip1*, *Arpc2/3*), lysosomal and autophagic trafficking and proteostasis (*Atg5*, *Gabarap*, *Ctsd*, *Vcp*), and intramembrane proteolysis and membrane protein catabolic processing (*Psen1*, *Adam10*). Although these genes are included in the SynGO database, they encode constitutive proteins; therefore, it is reasonable that they are highly expressed in microglia in both young and aged mice without microglial manipulation.

Collectively, these findings indicate that microglial repopulation in aged mice drives aberrant, age-specific derepression of neuronal gene expression.

### Microglial repopulation in aged mice impairs learning and cognitive flexibility

Given the aberrant gene expression programs of repopulated microglia in aged mice, we hypothesized that microglial repopulation might exacerbate age-related neural dysfunctions. To test this, we examined the effects of microglial repopulation on learning and cognitive flexibility, both of which decline with aging (Fig. 6A). Young and aged mice were subjected to a visual discrimination test to assess learning. We then switched the task rule by introducing a response direction test, which required mice to shift from a previously learned rule to a new one, thereby allowing us to evaluate attentional set shifting ability(Birrell & Brown, 2000; Higashida et al., 2018; H. Nagai, 2024). Without microglial manipulation, both young and aged mice learned the visual discrimination test to similar levels of accuracy, although learning was slower in aged mice (Fig. 6B-D). In the response direction test, aged mice performed worse than young mice, although some aged mice maintained cognitive flexibility comparable to that of young mice (Fig. 6E,F). These results indicate that aging impairs learning and cognitive flexibility with substantial individual variability. In contrast, after microglial repopulation, whereas young mice maintained their performance in visual discrimination learning, aged mice showed severe deficits: some aged mice never reached the behavioral criterion even when the number of trials per session was increased (Fig. 6B-D). Moreover, aged mice after repopulation failed to learn the response direction test with performance remaining near chance level, suggesting a loss of cognitive flexibility (Fig. 6E,F). Collectively, these findings demonstrate that microglial repopulation has age-specific deleterious effects on learning and cognitive flexibility.

**Figure 6.**
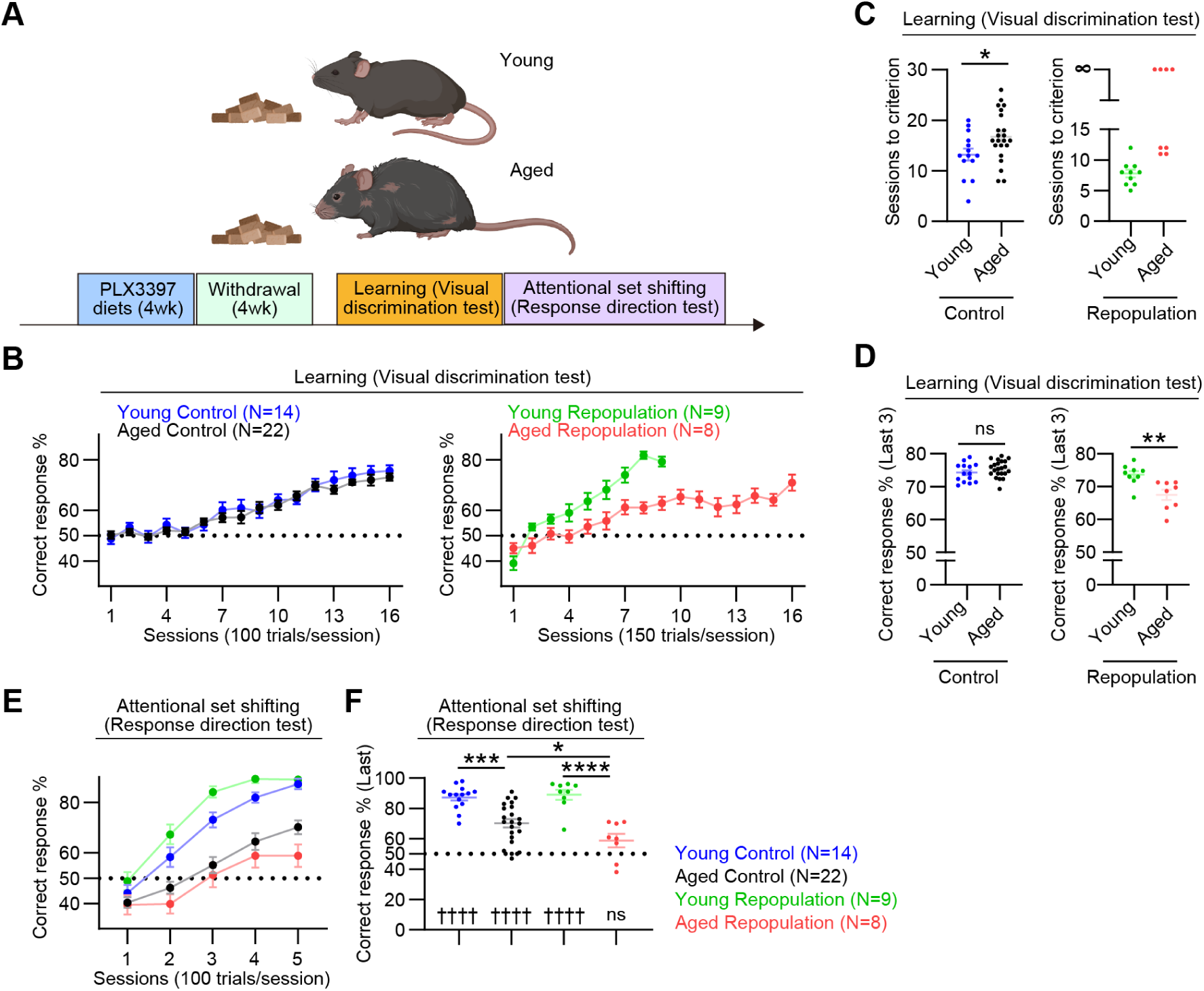
Microglial repopulation in aged mice impairs learning and cognitive flexibility. (A) Experimental schedule for behavioral tests. Young and aged mice were fed control diets (control) or PLX3397-containing diets for 4 weeks with an additional 4-week withdrawal period (repopulation). The mice then underwent visual discrimination and response direction tests to assess learning and attentional set shifting abilities. (B) Correct response rates in the visual discrimination test across sessions. (C) Number of sessions required to reach the performance criterion. (D) Correct response rates averaged across the last three sessions. (E) Correct response rates in the response direction test across sessions. (F) Correct response rate in the final session. Values are expressed as mean ± SEM. *p < 0.05, **p < 0.01, ***p < 0.001, ****p < 0.0001, ns; not significant for Student’s *t*-test (C, D) or for Holm-Sidak’s multiple comparisons test following one-way ANOVA (F). ††††p < 0.0001, ns; not significant for one-sample *t*-test comparing the distribution against the chance level (50%).

## 4. Discussion

In this study, we demonstrate that microglial repopulation after CSF1R inhibitor-mediated depletion has distinct and broad effects in aged brains compared with young brains, encompassing alterations in microglial gene expression states and cognitive function. In young mice, repopulated microglia largely maintain their pre-depletion gene expression profile and do not affect learning or cognitive flexibility. In contrast, in aged mice, repopulated microglia undergo a profound shift in gene expression patterns, characterized by a partial loss of expression of microglial identity genes, extensive reprogramming of immune-related gene expression, and derepression of neuronal gene expression. Concordant with these gene expression abnormalities, microglial repopulation severely impairs visual discrimination learning and attentional set shifting. Thus, although microglial repopulation has been regarded as beneficial to mitigate diverse neuropathologies, our findings indicate that aging can convert microglial repopulation into maladaptive reprogramming that ultimately exacerbates cognitive decline.

Because repopulated microglia in aged brains lose the expression of microglial identity genes, they are likely to acquire atypical functional characteristics. Among these genes, CX3CR1 is pivotal for maintaining microglial homeostasis. Indeed, CX3CR1 deficiency has been reported to impair synaptic refinement during postnatal development and to enhance microglial neurotoxicity across multiple neurodegenerative models(Cardona et al., 2006; Hoshiko, Arnoux, Avignone, Yamamoto, & Audinat, 2012). HexB (β-hexosaminidase B), whose expression is similarly lost in repopulated microglia in aged brains, encodes a lysosomal enzyme required for the degradation of the ganglioside GM2; its deficiency, as in Sandhoff disease, drives progressive neurodegeneration(Leal et al., 2020). Accordingly, loss of these genes could contribute to the exacerbation of cognitive decline associated with microglial repopulation after aging. Although microglia contribute to neuroinflammation in diverse contexts, including neurodegeneration, repopulated microglia in aged mice show reduced expression of innate and adaptive immune-related genes, as evidenced by decreased expression of genes encoding immune cell-surface receptors and of type I interferon-related genes, while also exhibiting gene expression changes indicative of enhanced type II interferon signaling. Thus, repopulated microglia could impair neuronal function through mechanisms distinct from those of inflammation-associated microglia.

A prominent and unexpected feature of repopulated microglia in aged mice is the robust induction of synapse-associated genes, including components of synaptic vesicle trafficking/release machinery and neurotransmitter receptor pathways. This observation raises the possibility that these aberrant gene expression programs enable microglia to interact with glutamatergic synaptic transmission that supports prefrontal-dependent cognitive function. It has been reported that, during postnatal development, a subset of microglia expressing GABA_B_ receptors selectively senses GABA, thereby contributing to inhibitory synapse refinement without affecting excitatory synapses(Favuzzi et al., 2021). Likewise, repopulated microglia in aged brains that acquire expression of glutamatergic receptors could selectively sense glutamate and modulate glutamatergic synaptic transmission. It remains to be determined whether and how microglia exert their effects on glutamatergic transmission, potentially via the secretion of effector molecules within the local synaptic milieu and the phagocytosis of synaptic components. Ectopic expression of genes related to synaptic vesicle trafficking and release machinery could alter the stimuli that trigger vesicle secretion from microglia and the effector molecules delivered by these secreted vesicles. Although direct evidence for a causal role of ectopic expression of synaptic and neuronal genes is still lacking, elucidating how atypical repopulated microglia in aged mice communicate with neurons will be key to understanding how microglial repopulation exacerbates age-related cognitive decline.

Whereas microglia-driven neuroinflammation has been proposed to underlie age-related cognitive decline, in this study, young and aged mice without microglial manipulation show only marginal differences in microglial gene expression patterns. In addition, microglial gene expression patterns do not exhibit notable individual variability among aged mice, unlike the variability observed in their performance on attentional set shifting. Thus, during physiological aging, cognitive decline and its individual variability could be driven by mechanisms other than microglial gene expression. Because resilient aged individuals are thought to retain adaptive mechanisms that preserve cognitive function, aberrant repopulated microglia could erode this adaptive capacity, thereby unmasking or accelerating cognitive decline.

In contrast to the deleterious effects of microglial repopulation observed in this study, beneficial effects have also been reported in various contexts(Colella et al., 2024; M. R. P. Elmore et al., 2018; Han et al., 2019; Hu et al., 2025; W. Wang et al., 2023). This apparent dichotomy suggests that repopulated microglia may consist of heterogenous populations that exert either beneficial or deleterious effects. The relative abundance of these populations may vary with multiple factors, including aging, brain region, and the duration and extent of microglial depletion, which may account for discrepancies across studies. For example, microglial repopulation has been reported to mitigate cognitive deficits in mouse models of neurodegenerative diseases such as Alzheimer’s disease; however, these effects were evaluated at relatively young ages across a limited set of behavioral domains, and under a certain depletion-repopulation condition(W. Wang et al., 2023). Thus, it remains unclear whether comparable benefits can be achieved in aged animals, when these diseases typically emerge. In addition, repletion and repopulation protocols need to be systematically optimized across behavioral domains.

Finally, how repopulated microglia acquire beneficial or deleterious properties remains an important open question. Because repopulated microglia derive from residual microglia that survive depletion(Huang et al., 2018), this residual pool may possess context-dependent properties that are beneficial in some settings and deleterious in others. Alternatively, residual microglia may be relatively homogeneous at baseline but acquire beneficial or harmful features during the repopulation process. Accordingly, in the aged brain, deleterious phenotypes after microglial repopulation could arise from cell-intrinsic impairments, such as senescence-associated epigenetic and metabolic derangements and/or from cell-extrinsic constraints imposed by an altered brain microenvironment, leading to distinct therapeutic implications. If dysfunction is primarily cell-intrinsic, replacing aged microglia with developmentally younger counterparts may be effective. By contrast, if maladaptive states are chiefly driven by extrinsic factors, replacement alone is unlikely to suffice without concomitant remodeling of the niche. Disentangling the relative contributions of cell-intrinsic and cell-extrinsic determinants will therefore be critical for evaluating whether microglial repopulation or replacement can be safely and effectively harnessed to treat age-related cognitive decline and neurodegenerative disorders.

## Acknowledgment

We thank Akemi Ando, Mikiko Suzuki, Misako Takizawa and Kaori Yamabe for secretarial help and Hiroko Iwamura for technical help. This study was supported in part by grants from AMED (JP25wm0625121 to H.N., JP24wm0425001, 25zf0127010, 25zf0127012, 25wm0625324 to T.F.), grants from JST Moonshot R&D (JPMJMS239F), Grant-in-Aid for Transformative Research Areas (23H04234 to T.F.) and Leading Initiative for Excellent Young Researchers (LEADER to H.N.) from the Ministry of Education, Culture, Sports, Science and Technology in Japan, Grants-in-Aid for Scientific Research (21H04812, 24K22086 to T.F., 20K07288, 23K06358 to H.N.) from the Japan Society for the Promotion of Science in Japan, and research grants from the Uehara Memorial Foundation (H.N.), Japan Foundation for Applied Enzymology (H.N.), the KANAE foundation for the promotion of medical science (H.N.), SENSHIN Medical Research Foundation (H.N., T.F.), Meiji Yasuda Life Foundation of Health and Welfare (H.N.), Daiichi Sankyo Foundation of Life Science (T.F.), SRF (T.F.), and the Kazato Foundation (H.N.).

## Author contributions

R.Y., H.N., and T.F. designed the study; R.Y., H.N., Y.Z., C.N., M.T., and T.F. performed and analyzed the results; H.N. and T.F. wrote the manuscript.

## Data availability

The datasets generated during and/or analyzed during the current study are available from the corresponding author on reasonable request.

